# Trophoblast ferroptosis restricts SARS-CoV-2 spread in the placenta

**DOI:** 10.64898/2026.01.26.701742

**Authors:** Eliza R McColl, Deepak Kumar, Brittany R Jones, Nikole Marks, Emily Diveley, Rowan Karvas, Thorold W Theunissen, Jeannie C Kelly, Indira U Mysorekar

## Abstract

Prenatal SARS-CoV-2 infection is associated with adverse pregnancy outcomes, but placental mechanisms that restrict viral spread remain unclear. Here we show that SARS-CoV-2 exposure induces ferroptosis-linked iron dysregulation in the placenta as a host defense. Human placentas from early gestation SARS-CoV-2–exposed pregnancies exhibited persistent viral protein expression at term, iron accumulation, disrupted localization of iron transport proteins, and reduced expression of the ferroptosis inhibitor, GPX4. In trophoblast cells and newly generated stem cell-derived trophoblast organoids (SC-TOs) with physiological apical-out polarity, infection with live SARS-CoV-2 Delta variant suppressed expression of iron efflux transporter, ferroportin and ferroptosis inhibitors, GPX4 and PLA2G6, promoting lipid peroxidation and ferroptotic signaling. Sub-lethal pharmacological activation of ferroptosis reduced viral titers in trophoblasts, indicating an antiviral function. Together, these results uncover a new mechanism through which the placenta attempts to restrict SARS-CoV-2 replication. However, this protective response is accompanied by placental iron sequestration, which may compromise maternal-fetal iron transfer and help explain iron deficiency and anemia reported in infants born after prenatal SARS-CoV-2 exposure, highlighting a delicate balance between iron and ferroptosis-mediated protection and damage with implications for pregnancy outcomes.

## Introduction

Since its emergence in 2019, SARS-CoV-2 has claimed the lives of over 7 million people.^1^ Pregnant people are more vulnerable to increased COVID-19 disease severity and morbidity than their non-pregnant counterparts.^2–4^ Infection with SARS-CoV-2 also increases chances of adverse pregnancy outcomes including preterm birth, stillbirth, and preeclampsia.^5,6^ Furthermore, longitudinal studies have revealed long-term impacts of SARS-CoV-2 infection during pregnancy on offspring neurodevelopment.^7–9^ While vaccination has significantly reduced transmission and mortality in the general public, vaccine uptake in pregnant individuals is lower than in the non-pregnant population,^10–13^ leaving them at increased risk for severe infection and adverse outcomes. However, the mechanisms underlying infection-mediated adverse pregnancy outcomes have not been fully characterized.

The placenta, which expresses the SARS-CoV-2 entry receptor angiotensin-converting enzyme 2 (ACE2), is susceptible to SARS-CoV-2 infection.^14,15^ However, instances of transplacental transmission to the fetus are rare,^16^ potentially due to undefined mechanisms that limit SARS-CoV-2 replication in placental cells.^17–20^ Despite limited viral replication and vertical transmission, SARS-CoV-2 causes a number of changes in placental function and pathology including SARS-CoV-2 placentitis (chronic histiocytic intervillositis, perivillous fibrin deposition, and trophoblast necrosis),^21^ DNA damage,^22,23^ increased syncytial knot formation,^24,25^ dysregulated renin-angiotensin signaling,^15,24^ and impaired autophagic flux.^26^ Several studies have also described increased oxidative stress in placentas of SARS-CoV-2-infected individuals.^22,23,27,28^ This suggests that SARS-CoV-2 infection could trigger oxidative stress-induced cell death, such as ferroptosis, an iron-dependent form of cell death characterized by reduced redox response and increased lipid peroxidation.^29^ Emerging evidence supports a role for ferroptosis in various pregnancy complications,^30,31^ particularly preeclampsia.^32–35^ Thus, infection-mediated activation of placental ferroptosis could be a mechanism through which COVID-19 leads to adverse pregnancy outcomes. SARS-CoV-2 has been shown to activate or increase susceptibility to ferroptosis in the heart,^36^ lungs,^37,38^ and plasma.^39,40^ Dysregulated iron homeostasis, particularly hyperferritinemia, is also a common finding in COVID-19 patients.^41^ Moreover, ferroptosis and iron dysregulation have been implicated in the development and prognosis of long COVID.^39,41,42^ However, whether SARS-CoV-2 similarly affects iron homeostasis and ferroptosis in the placenta and the resulting implications of such changes for host defense mechanisms or adverse pregnancy outcomes remain unclear.

Given their ability to recapitulate multiple trophoblast subtypes in 3-dimensional configurations, trophoblast organoids are a powerful *in vitro* model for studying the placenta.^43^ However, traditional trophoblast organoid models exhibit inverse polarity, with an inner core of syncytiotrophoblasts (STB) surrounded by an outer layer of cytotrophoblasts (CTB), termed STB^IN^. This reverse orientation of trophoblast layers poses challenges for modelling directional processes such as placental nutrient transport and vertical transmission of pathogens. Indeed, we^44^ and others^45^ have found that STB^IN^ organoids show very limited susceptibility to infection with SARS-CoV-2, likely due to the fact that ACE2 is more highly expressed in the STB layer^15,45^ and therefore inaccessible to the virus in the STB^IN^ conformation. Recently, several groups have reported methods to generate trophoblast organoid models with an outer STB layer surrounding an inner CTB core, termed STB^OUT^.^46–48^ In addition to more closely resembling the *in vivo* orientation of trophoblast layers in the placenta, STBs from STB^OUT^ organoids exhibit enhanced gene expression of hormone transcripts and various nutrient transport processes,^49^ suggesting closer resemblance to *in vivo* STB function. Moreover, exposing the STB layer to the culture environment should more closely mimic susceptibility to pathogens transmitted vertically from maternal blood. However, existing methods for generating STB^OUT^ require additional considerations such as culture agitation,^47^ seeding with collagen-coated beads,^46^ or growth in custom-fabricated culture plates,^48^ adding additional technical hurdles.

Here, we present a comprehensive evaluation of ferroptosis in the placenta due to SARS-CoV-2 infection. Using human placental tissue from individuals infected during pregnancy, we show that SARS-CoV-2 exposure is associated with iron accumulation, altered localization of iron transport and storage proteins, and reduced expression of the ferroptosis inhibitor glutathione peroxidase 4 (GPX4) in the placenta. Through live infection of two *in vitro* trophoblast models, including newly developed 3D stem cell-derived trophoblast organoids (SC-TOs) with an outer STB layer, we further demonstrate that active SARS-CoV-2 infection impairs trophoblast iron efflux and activates ferroptosis signaling. Pharmacological activation of ferroptosis signaling reduced viral titers in trophoblasts, suggesting that activation of ferroptosis may be a protective mechanism to restrict viral replication in the placenta and limit transplacental transmission. Together, these findings identify ferroptosis as a previously unrecognized placental response to SARS-CoV-2 infection, while indicating that sustained activation of this pathway may compromise placental function and contribute to adverse pregnancy outcomes associated with COVID-19.

## Results

### Patient demographics

Clinical characteristics of patient samples used in this study are listed in **Table 1**. Maternal age, race, comorbidities, and perinatal outcomes did not differ significantly between groups. Of those positive for SARS-CoV-2, 80% were infected during the first or second trimesters. Infections mainly occurred during the Omicron variant period (n=8) followed by the Delta variant period (n=2). While 70% of infected patients were symptomatic, none were hospitalized, administered oxygen therapy, or received breathing support during infection.

**Table 1:**
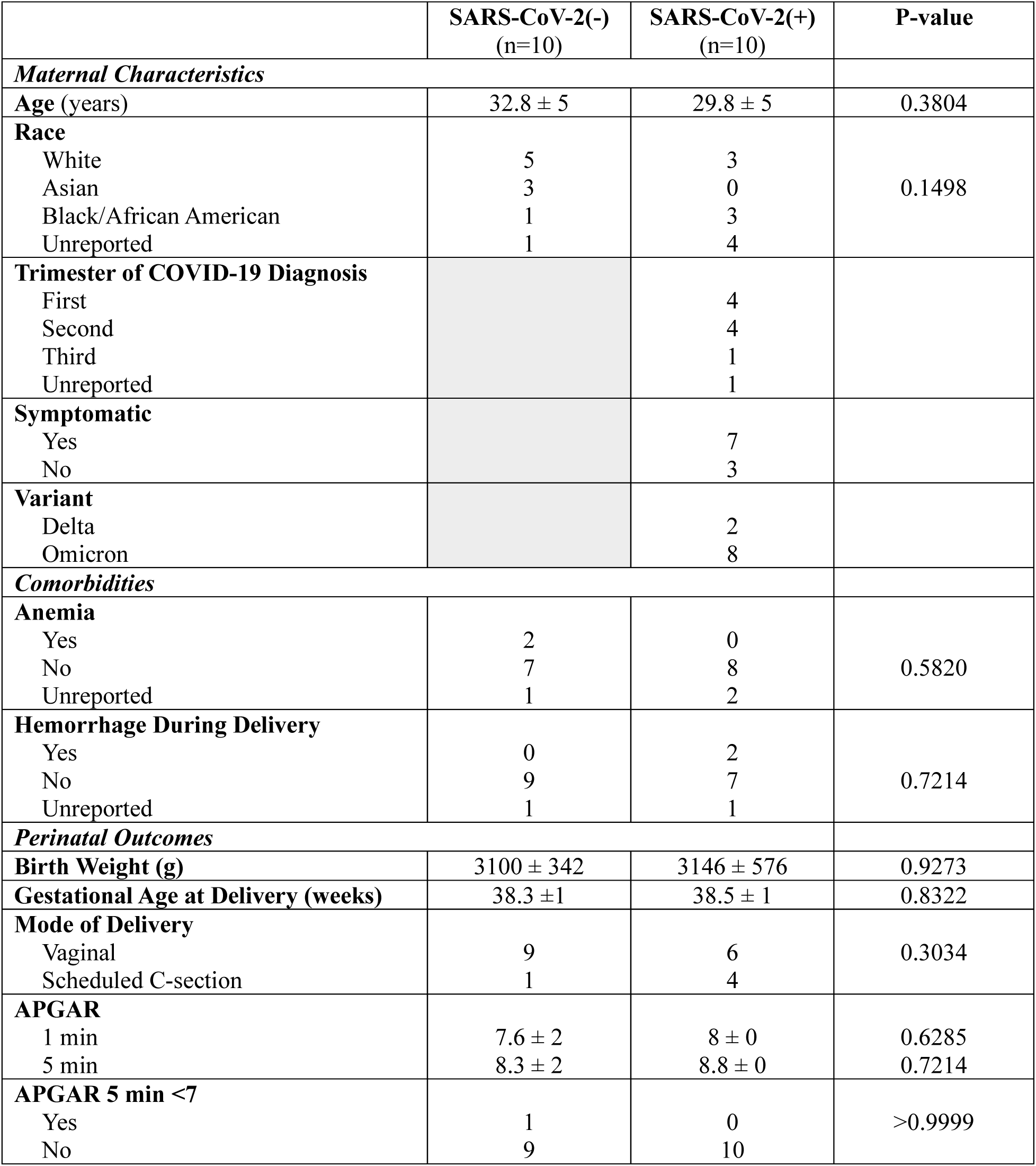
Demographic information for patient samples. Comparison of maternal characteristics, comorbitidites, and perinatal outcomes for patient samples used in this study. P-values were obtained using Mann-Whitney U-tests for continuous variables and Fisher’s Exact test for categorical variables.

|  | SARS-CoV-2(-)<br>(n=10) | SARS-CoV-2(+)<br>(n=10) | P-value |
| --- | --- | --- | --- |
| Maternal Characteristics |  |  |  |
| Age (years) | 32.8 ± 5 | 29.8 ± 5 | 0.3804 |
| Race |  |  | 0.1498 |
| White | 5 | 3 |  |
| Asian | 3 | 0 |  |
| Black/African American | 1 | 3 |  |
| Unreported | 1 | 4 |  |
| Trimester of COVID-19 Diagnosis |  |  |  |
| First |  | 4 |  |
| Second |  | 4 |  |
| Third |  | 1 |  |
| Unreported |  | 1 |  |
| Symptomatic |  |  |  |
| Yes |  | 7 |  |
| No |  | 3 |  |
| Variant |  |  |  |
| Delta |  | 2 |  |
| Omicron |  | 8 |  |
| Comorbidities |  |  |  |
| Anemia |  |  | 0.5820 |
| Yes | 2 | 0 |  |
| No | 7 | 8 |  |
| Unreported | 1 | 2 |  |
| Hemorrhage During Delivery |  |  | 0.7214 |
| Yes | 0 | 2 |  |
| No | 9 | 7 |  |
| Unreported | 1 | 1 |  |
| Perinatal Outcomes |  |  |  |
| Birth Weight (g) | 3100 ± 342 | 3146 ± 576 | 0.9273 |
| Gestational Age at Delivery (weeks) | 38.3 ±1 | 38.5 ± 1 | 0.8322 |
| Mode of Delivery |  |  | 0.3034 |
| Vaginal | 9 | 6 |  |
| Scheduled C-section | 1 | 4 |  |
| APGAR |  |  |  |
| 1 min | 7.6 ± 2 | 8 ± 0 | 0.6285 |
| 5 min | 8.3 ± 2 | 8.8 ± 0 | 0.7214 |
| APGAR 5 min <7 |  |  | >0.9999 |
| Yes | 1 | 0 |  |
| No | 9 | 10 |  |

### Placentas from SARS-CoV-2-exposed pregnancies exhibit persistent viral protein expression and iron accumulation

Distinct patches of villi containing SARS-CoV-2 Spike protein were observed in patients from the SARS-CoV-2(+) cohort (**Figure 1A**). Spike expression was restricted to the STB layer. Importantly, active infection at time of collection was not required for Spike protein staining. **Figure 1A** represents an individual infected during the first trimester, who still displayed Spike protein staining when the placenta was collected at term.

**Figure 1:**
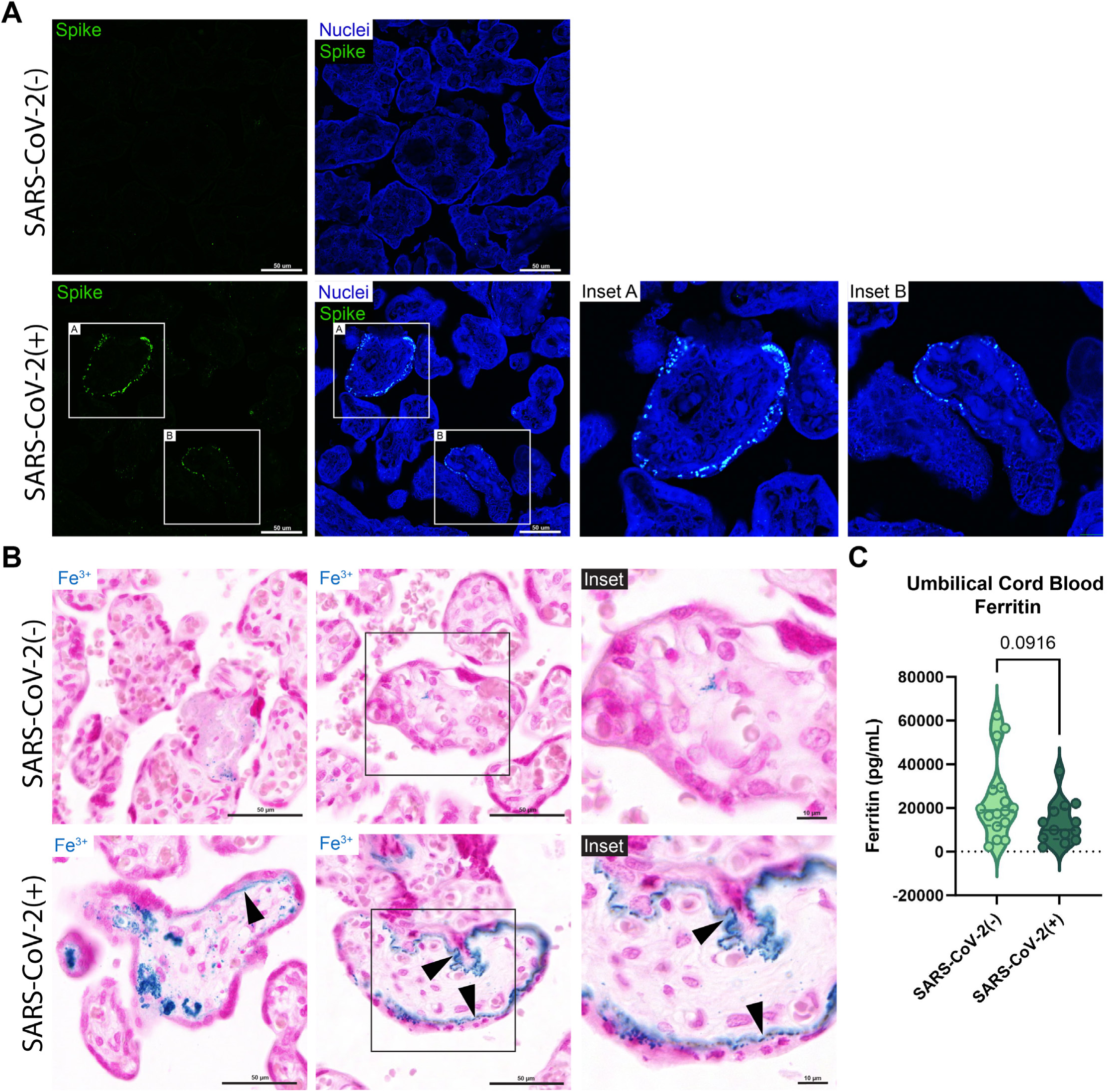
SARS-CoV-2 infection is associated with persistent viral protein expression and placental iron accumulation. **(A)** Representative immunofluorescence images showing expression of the SARS-CoV-2 spike protein in human placental tissue samples. Image shown is from a patient infected during the first trimester. **(B)** Prussian blue staining of human placental sections, with ferric iron stained blue. Representative images are from two different patient samples per group. Black arrows indicate linear iron deposits along the basolateral syncytiotrophoblast membrane. **(C)** Ferritin levels in umbilical cord blood from uninfected and infected pregnancies, as measured by Luminex. Significance was determined via Mann-Whitney U-test.

Iron accumulation in placental tissue was evaluated via Prussian Blue staining. In contrast to SARS-CoV-2(-) placentas, which showed areas of diffuse iron staining in patches of fibrin or in villous stroma, SARS-CoV-2(+) placentas exhibited strong Prussian Blue staining indicating accumulation of iron (**Figure 1B**). Iron deposits in SARS-CoV-2(+) placentas were present in patches of villi and often localized to the basolateral membrane of the STB. Consistent with placental iron accumulation, umbilical cord blood from SARS-CoV-2(+) pregnancies exhibited a trend towards decreased ferritin levels (**Figure 1C**), suggesting a lack of iron being transferred to the fetus. This could contribute to the increased incidences of anemia reported in neonates exposed to SARS-CoV-2 in utero.^50,51^

### SARS-CoV-2-exposed placentas show altered localization of iron transport and storage proteins

We further examined whether SARS-CoV-2 infection is associated with changes in placental iron transport and storage. Iron from maternal blood is taken into the placenta via the transferrin receptor (TFRC), stored bound to ferritin (FTH), and effluxed towards fetal circulation through ferroportin (FPN) (**Figure 2A**). Localization and expression of TFRC were unaffected by SARS-CoV-2 exposure (**Figure 2B,C**). While overall levels of FTH were not significantly altered (**Figure 2E**), localization of FTH showed distinct differences between SARS-CoV-2(-) and SARS-CoV-2(+) placentas. In unexposed placentas, FTH was distributed diffusely throughout the stroma (**Figure 2D**). In contrast, patches of SARS-CoV-2(+) placentas exhibited strong FTH expression within particular cells of the stroma, further supporting our hypothesis that iron is sequestered by SARS-CoV-2(+) placentas. Similarly, while levels of FPN were not significantly altered (**Figure 2G**), a drastic change in localization was observed. To perform efflux of iron from the placenta to fetal circulation, FPN is typically localized to the basolateral membrane of the STB, as shown in SARS-CoV-2(-) placentas (**Figure 2F**). However, SARS-CoV-2(+) placentas exhibited expression of FPN on both the basolateral and the maternal-facing microvillous membrane, potentially reflecting an attempt to transport excess iron back into maternal circulation. Together with increased Prussian Blue iron staining, these results suggest that SARS-CoV-2 exposure during pregnancy is associated with disruptions in iron transport and accumulation of iron in the placenta that persists throughout pregnancy, even after active infection has ceased.

**Figure 2:**
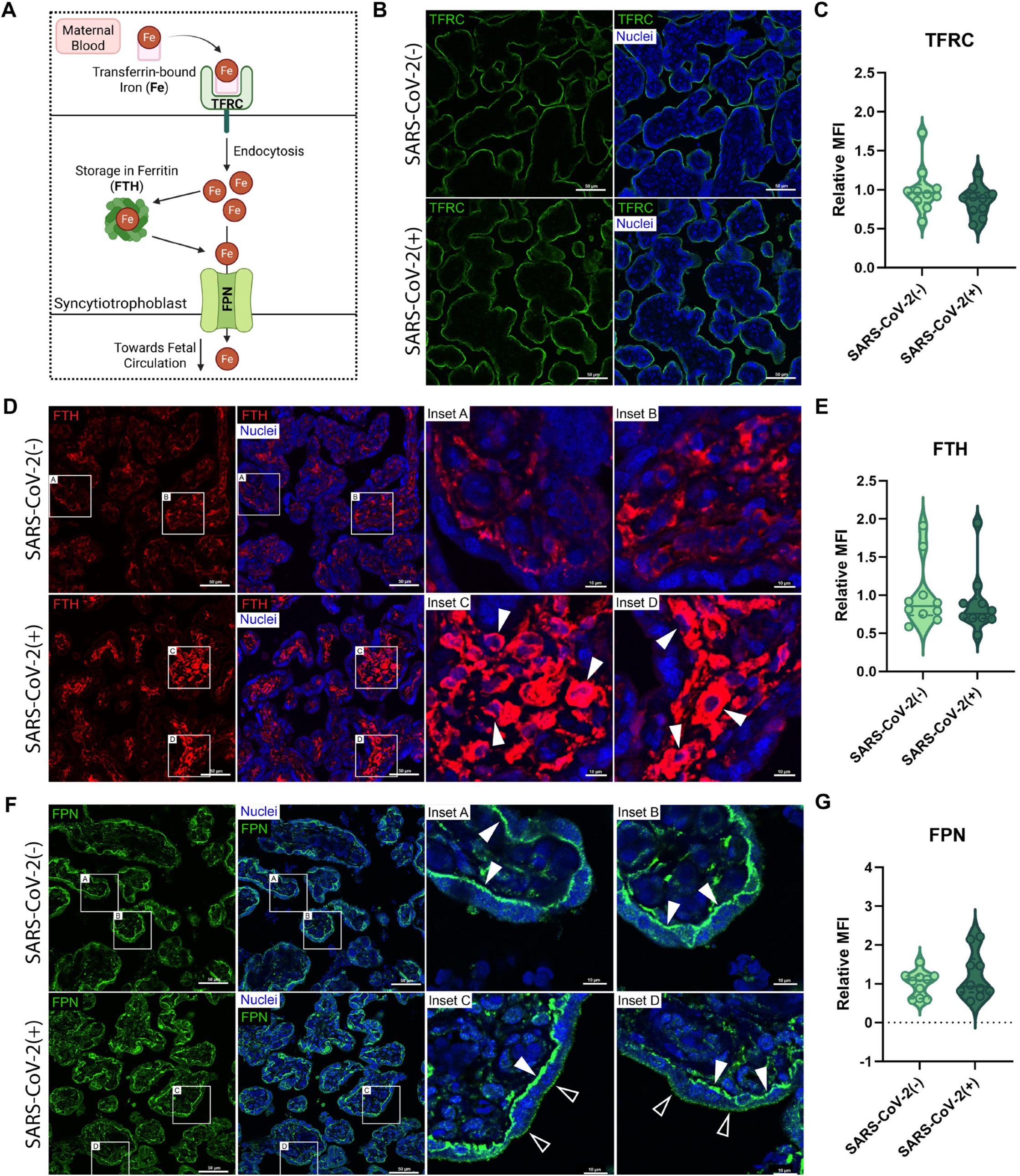
SARS-CoV-2 infection disrupts localization of proteins involved in placental iron transport and storage. **(A)** Schematic depicting placental uptake of transferrin-bound iron via TFRC, storage in ferritin (FTH), and efflux across the basolateral membrane via FPN. **(B)** Immunofluorescence images depicting TFRC localization (green) in human placental samples, as quantified in **(C)**. **(D)** Immunofluorescence images of FTH localization (red) in human placental samples. White arrows in insets point to cells with FTH accumulation. **(E)** Quantification of FTH fluorescence. **(F)** Immunofluorescence images of FPN localization (green) in human placental samples. White-filled arrows show canonical basolateral syncytiotrophoblast localization, while white-outlined arrows show abnormal localization on microvillous membrane. **(G)** Quantification of fluorescent intensity for FPN. Fluorescence images (B,D,F) are representative of n=10 patient samples per group. Quantifications of fluorescent intensity (C,E,G) were compared with Mann-Whitney U-tests, with each point corresponding to the average MFI of five images taken per slide, with each slide (n=10 per group) corresponding to a different patient.

### SARS-CoV-2-exposed placentas exhibit decreased expression of the ferroptosis inhibitor GPX4

To determine whether the observed iron accumulation may be associated with activation of ferroptosis, we examined protein and biochemical markers of ferroptosis in exposed placentas. Consistent with activation of ferroptosis, the expression of the ferroptosis inhibitor GPX4 was significantly decreased in SARS-CoV-2(+) placentas (**Figure 3A,B**). However, no significant changes in either total or reduced glutathione levels were observed (**Figure 3C,D**). Several markers of lipid peroxidation were examined. In contrast to previous reports,^52,53^ acyl-CoA synthetase long-chain family member 4 (ACSL4), a protein upstream of lipid peroxidation, showed a strong trend towards decreased expression (**Figure 3E,F**). Despite reduced GPX4 expression and iron accumulation, SARS-CoV-2(+) placentas showed no significant change in two markers of lipid peroxidation: malondialdehyde (MDA) levels determined by TBARS assay, or 4-hydroxynoneal (4HNE) levels measured by western blotting (**Figure 3G,H**). Together, this suggests that while SARS-CoV-2(+)-exposed placentas show some signs of ferroptosis, active lipid peroxidation may not persist after active infection has ceased.

**Figure 3:**
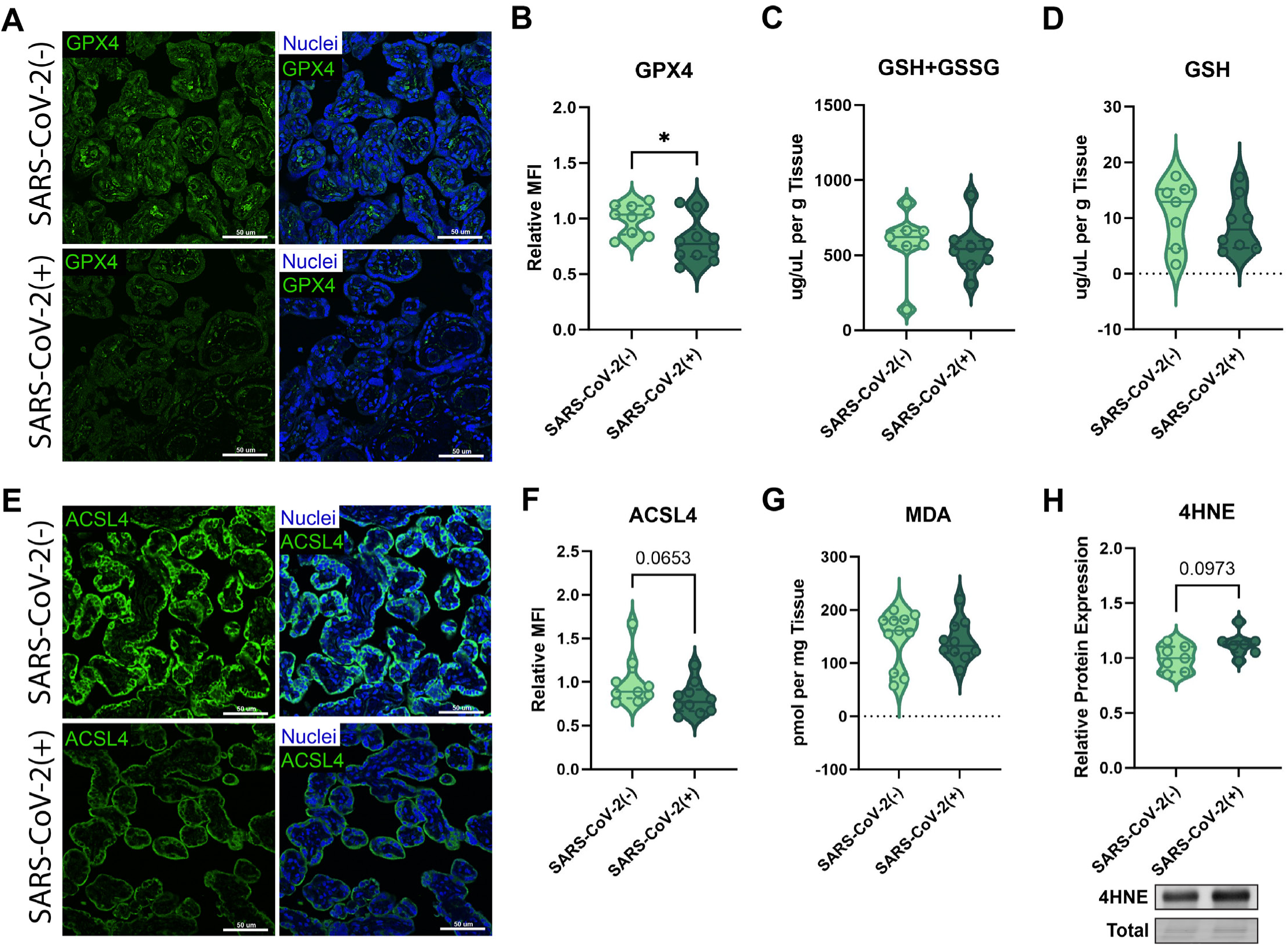
SARS-CoV-2 infection decreases GPX4 expression in the placenta. **(A)** Representative immunofluorescence image showing GPX4 expression (green). **(B)** Quantification of GPX4 fluorescent intensity. **(C)** Total and **(D)** reduced glutathione levels in human placental tissue (n=7-10) quantified via colorimetric assay. **(E)** Representative immunofluorescence image showing ACSL4 expression (green). **(F)** Quantification of ACSL4 fluorescent intensity. **(G)** MDA (lipid peroxidation marker) levels in human placental tissue quantified via TBARS assay (n=7-10). **(H)** Expression of the lipid peroxidation marker 4HNE in human placental tissue determined via Western blotting (n=7). Fluorescence images (A,E) are representative of n=10 patient samples per group. For quantification of fluorescent intensity (B,F) each point represents the average MFI of five images taken per slide, with each slide (n=10 per group) corresponding to a different patient. Solid lines on violin plots signify the median, with dotted lines representing upper and lower quartiles. Significance was determined using Mann-Whitney U-tests (* p<0.05).

### SARS-CoV-2 Delta variant disrupts iron transport and activates ferroptosis in trophoblast cells

The human placentas examined in this study were from individuals infected with SARS-CoV-2 at any point during pregnancy, and the majority of our cohort was infected during the first or second trimester. As such, placentas collected and examined at term may not reflect changes that occur during active infection. To determine whether altered iron transport, activation of ferroptosis, and increased lipid peroxidation occur during active infection, we infected JEG-3 trophoblast cells with the SARS-CoV-2 Delta variant (**Figure 4A**). The Delta variant was chosen due to its strong association with adverse pregnancy outcomes.^54,55^ While JEG-3 cells exhibit low levels of SARS-CoV-2 infection, we have shown that they sustain molecular and biochemical changes consistent with other models of SARS-CoV-2 infection in the placenta.^26^ Indeed, infection of JEG-3 cells caused a significant and robust decrease in FPN expression after 48 hours post-infection (hpi) (**Figure 4C**). In contrast, levels of the iron uptake protein TFRC (**Figure 4B**) and iron storage proteins FTH (**Figure 4D**) and FTL (**Figure 4E**) were not significantly affected. This suggests that the SARS-CoV-2 Delta variant causes reduced iron efflux, resulting in trophoblast iron accumulation. It is possible that the reduction of FPN levels is due to an observed 10-fold increase in iron-responsive element-binding protein 2 (IRP2) expression in infected cells (**Figure 4H**), as increased IRP activity represses translation of *SLC40A1*, the gene encoding FPN, leading to reduced iron efflux (**Figure 4F**).^56^ IRP induction also limits FTH translation while stabilizing TFRC expression, thus facilitating iron uptake and increasing the labile iron pool,^56^ a known trigger of ferroptosis. Since intracellular iron accumulation has been shown to enhance SARS-CoV-2 infection in other cell types,^57,58^ we next examined whether iron loading with ferric ammonium citrate (FAC) influenced viral replication in JEG-3 cells. However, 48 hours of treatment with FAC had no significant impact on viral titers (**Figure 4I**).

**Figure 4:**
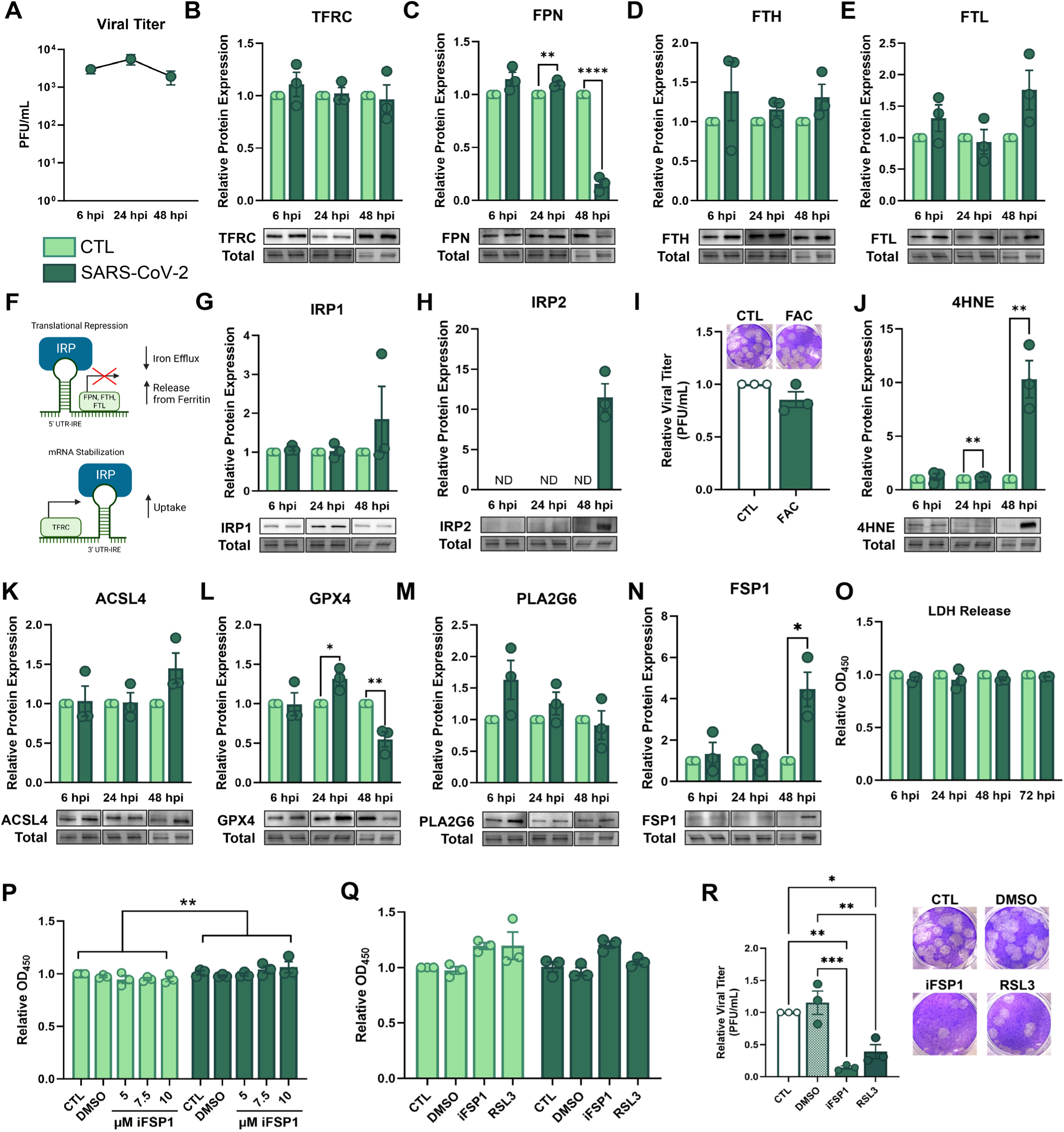
SARS-CoV-2 Delta variant reduces iron efflux and activates ferroptosis in cytotrophoblasts. **(A)** Viral titers of SARS-CoV-2 Delta variant in supernatants from JEG-3 cells infected over 48 hours with an MOI of 1. Titers were determined via plaque assay. **(B-E)** Protein expression of iron transport and storage proteins in SARS-CoV-2-infected JEG-3 cells at 6, 24, and 48 hpi, as determined by western blotting. **(F)** Schematic of regulation of iron transport and storage expression by iron regulatory proteins (IRPs). **(G,H)** Protein expression of IRPs in infected JEG-3 cells, as determined by western blotting. **(I)** Impact of excess iron supplementation (FAC) on viral titers in JEG-3 cells, as determined via plaque assay. **(J-N)** Protein expression of ferroptosis markers and inhibitors in SARS-CoV-2-infected JEG-3 cells over course of infection. Expression was measured by western blotting. **(O)** Cytotoxicity of SARS-CoV-2 Delta variant in JEG-3 cells as measured by LDH release over course of infection. **(P)** Cytotoxicity of JEG-3 cells treated with increasing concentrations of the FSP1 inhibitor, iFSP1, with or without infection. Cytotoxicity was determined via LDH assay. **(Q)** Cytotoxicity of small molecules iFSP1 and RSL3 in JEG-3 cells with or without SARS-CoV-2 infection, as measured by LDH release. **(R)** Viral titers in supernatants from JEG-3 cells infected with SARS-CoV-2 for 48 hours, following pre-treatment and post-inoculation exposure to iFPS1 and RSL3. Viral titers were determined via plaque assay. Data represents average of n=3 independent replicates per group ± SEM. Significance determined by Student’s unpaired t-test except for P and Q which were determined by two-way ANOVA and R which was determined by one-way ANOVA. For all tests, *p<0.05, **p<0.01, ***p<0.001, ****p<<0.001.

Next, we examined the impact of active infection on ferroptosis markers. Consistent with ferroptosis activation, SARS-CoV-2-infected cells showed a significant increase in the lipid peroxidation marker 4HNE starting at 24 hpi and reaching a 10-fold increase by 48 hpi (**Figure 4J**). Expression of ACSL4, while not significantly altered, showed a similar trend towards increased expression at 48 hpi (**Figure 4K**). To gain further insight into the mechanisms through which ferroptosis is activated during infection, we also examined upstream three inhibitors of ferroptosis: GPX4, phospholipase A2 group VI (PLA2G6), and ferroptosis suppressor protein 1 (FSP1).^59,60^ Consistent with our observations in human placentas, GPX4 protein expression was significantly decreased at 48 hpi (**Figure 4L**). Together with the observed reduction of FPN and increase in lipid peroxidation, this indicates that the SARS-CoV-2 Delta variant activates ferroptosis signaling in cytotrophoblasts. Interestingly, GPX4 was significantly increased at 24 hpi, suggesting a potential attempt to prevent ferroptosis activation early in infection. Similarly, PLA2G6 expression showed a trend towards increased expression at earlier timepoints followed by waning expression throughout infection, further supporting an early attempt to limit ferroptosis (**Figure 4M**). Finally, expression of FSP1 was substantially and significantly increased at 48 hpi (**Figure 4N**), signifying a potential compensatory mechanism against completion of ferroptosis in the event of reduced GPX4. Consistent with the notion of a compensatory mechanism, SARS-CoV-2-infected cells did not exhibit increased lactate dehydrogenase (LDH) release over 72 hours of infection (**Figure 4O**), suggesting a lack of cytotoxicity despite activated ferroptosis signaling.

To determine whether induction of FSP1 prevented infection-mediated cell death, we infected JEG-3 cells with SARS-CoV-2 in the presence of iFSP1, a potent FSP1 inhibitor.^61^ While infected and uninfected cells responded to increasing concentrations of iFSP1 differently (p < 0.01 column factor in two-way ANOVA), iFSP1 did not significantly induce cell death in infected cells (**Figure 4P**). Thus, the lack of ferroptosis-mediated cytotoxicity in SARS-CoV-2-infected JEG-3 cells appears to be due to an alternative mechanism. Despite no significant impact on cell death (**Figure 4Q**), inhibition of FSP1 significantly reduced viral titers in infected cells (**Figure 4R**). Similarly, sub-lethal treatment with the ferroptosis activator RSL3 significantly reduced viral titers. Together, this suggests that activation of ferroptosis signaling in trophoblasts may play a protective role by limiting viral dissemination.

### Culturing SC-TOs in suspension generates organoids with STB^OUT^ polarity that express ACE2 on their outer surface

We next sought to translate our findings from the JEG-3 model to a more physiologically complex 3D model using trophoblast organoids. Like most trophoblast organoid models, human trophoblast stem cell (hTSC)-derived SC-TOs grown in Matrigel exhibit inside-out polarity with an inner syndecan-1 (SDC-1)-positive STB core surrounded by an outer layer of E-cadherin (E-cad)-positive CTBs (**Figure 5A,B**), termed STB^IN^. STB^IN^ SC-TOs show very limited susceptibility to infection with SARS-CoV-2,^44,45^ likely due to an inability to access the inner ACE2-expressing STB layer.^15,45^. Thus, we developed a method for generating SC-TOs with an inner CTB core surrounded by an outer STB layer (STB^OUT^), which reflects the orientation of these cell layers *in vivo*. After initial seeding of hTSCs in Matrigel, SC-TOs were removed from Matrigel domes on Day 3 and cultured in suspension in AggreWell 400 plates (**Figure 5A**). Changing to suspension culture resulted in mature SC-TOs with STB^OUT^ polarity (**Figure 5B**). Of note, culturing SC-TOs in commercially-available AggreWell plates negates the need for seeding on collagen-coated beads or continuous agitation of suspension cultures, a methodological improvement over previous STB^OUT^ models.^46–48^ STB^OUT^ organoids generated in this manner begin to exhibit STB^OUT^ polarity as early as 24 hours after transfer to suspension culture, progressively increasing the degree of outer STB coverage over time (**Figure 5C**). By Day 6, over 90% of organoids exhibit some degree of outer STB coverage, with the majority being more than half covered with STB (**Figure 5D**). As such, subsequent infections were performed on culture Day 6. STB^OUT^ SC-TOs exhibited ACE2 expression on the outer membrane, suggesting they may be more susceptible to SARS-CoV-2 infection (**Figure 5E**). They also expressed proteins required for uptake (TFRC) and efflux (FPN) of iron, making them a viable model to study infection-mediated changes in iron transport and ferroptosis (**Figure 5F**).

**Figure 5:**
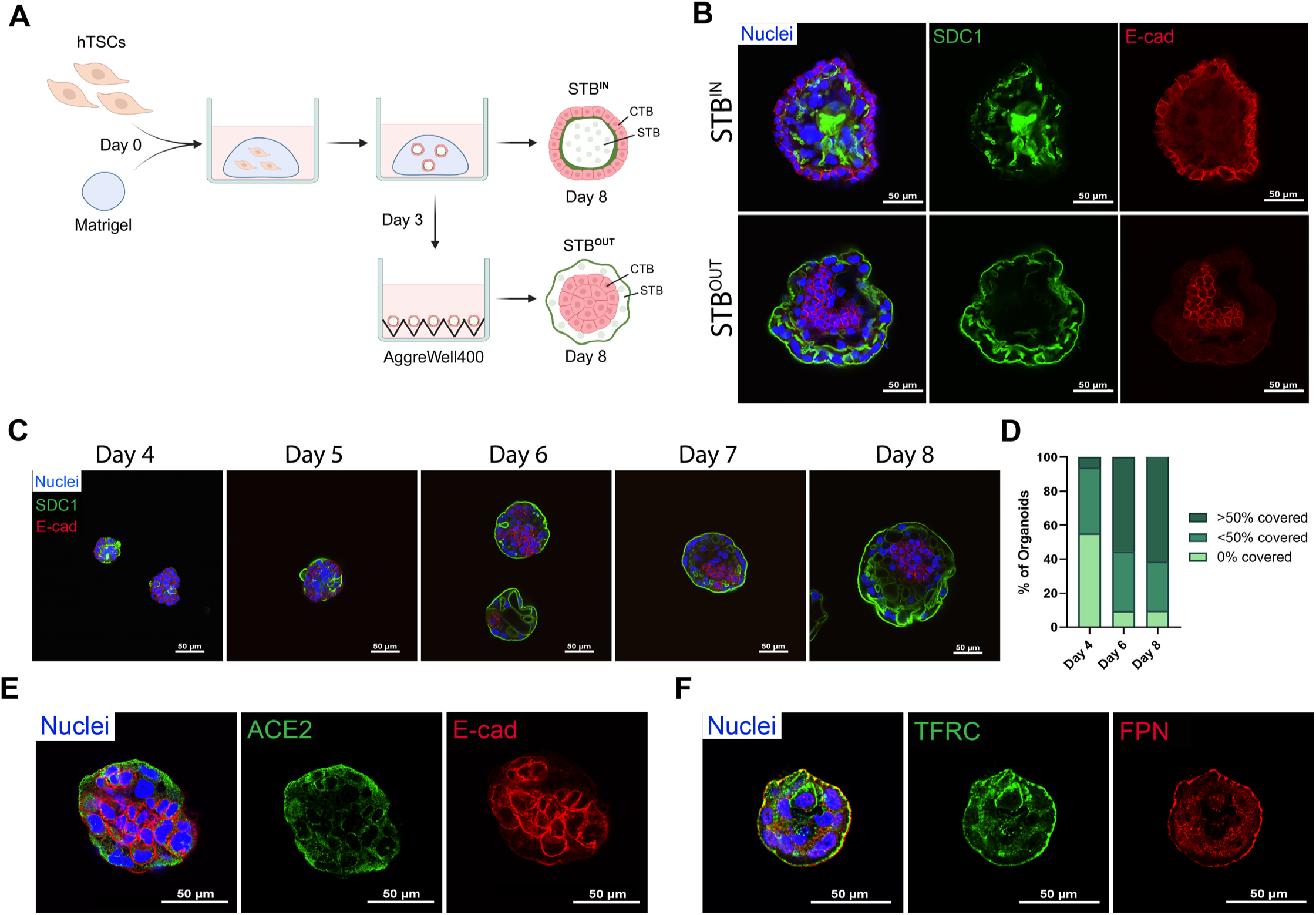
SC-TOs grown in suspension display apical-out polarity (STB^OUT^) with an outer STB layer expressing ACE2 and iron transport proteins. **(A)** Schematic showing methods for cultivating STB^IN^ versus STB^OUT^ SC-TOs. **(B)** STB^IN^ versus STB^OUT^ SC-TOs on Day 8 of culture. STB^OUT^ SC-TOs exhibit an outer STB layer marked with SDC1 (green) surrounding an inner CTB core marked with E-cad (red). **(C)** Time course showing growth and progression of STB outer coverage as marked with STB marker SDC1 (green) versus internal CTB marker E-cad (red). **(D)** Percentage of SC-TOs exhibition greater than or less than 50% outer STB coverage throughout culture period. Data represents average from three independent replicates, quantified as described in methods. **(E)** Representative image of Day 6 STB^OUT^ SC-TOs exhibiting ACE2 localization (green) to the outer STB layer. The inner CTB core is marked with E-cad (red). **(F)** Representative image of Day 6 STB^OUT^ SC-TOs expressing iron transport proteins TFRC (green) and FPN (red) on the outer STB surface. All images are representative of n=3 independent replicates.

### SARS-CoV-2 Delta variant impairs iron efflux and activates ferroptosis in STB^OUT^ trophoblast organoids

Consistent with localization of ACE2 to the outer STB layer, STB^OUT^ organoids were susceptible to infection with the SARS-CoV-2 Delta variant (**Figure 6A, B**). However, viral titers decreased over time, which aligns with previous suggestions that the placenta exhibits mechanisms to restrict viral replication.^17–20^ Of note, viral protein expression was restricted almost exclusively to the STB layer (**Figure 6B**), further supporting previous reports that STBs are more permissive to SARS-CoV-2 infection than CTBs.^45,62,63^ STB^OUT^ SC-TOs exhibited significantly increased LDH release at 24 and 48 hpi, consistent with the notion that SARS-CoV-2 infection with the Delta variant induces trophoblast cell death. Similarly to JEG-3 cells, SARS-CoV-2 Delta variant significantly reduced FPN expression in STB^OUT^ organoids at 72 hpi, whereas TFRC, FTH, and FTL were not significantly altered (**Figure 6D-G**). Expression of IRP1 and IRP2 was not significantly altered at 72 hpi (**Figure 6H, I**). Together, this supports our observations that SARS-CoV-2 Delta variant reduces iron efflux, potentially leading to trophoblast iron accumulation.

**Figure 6:**
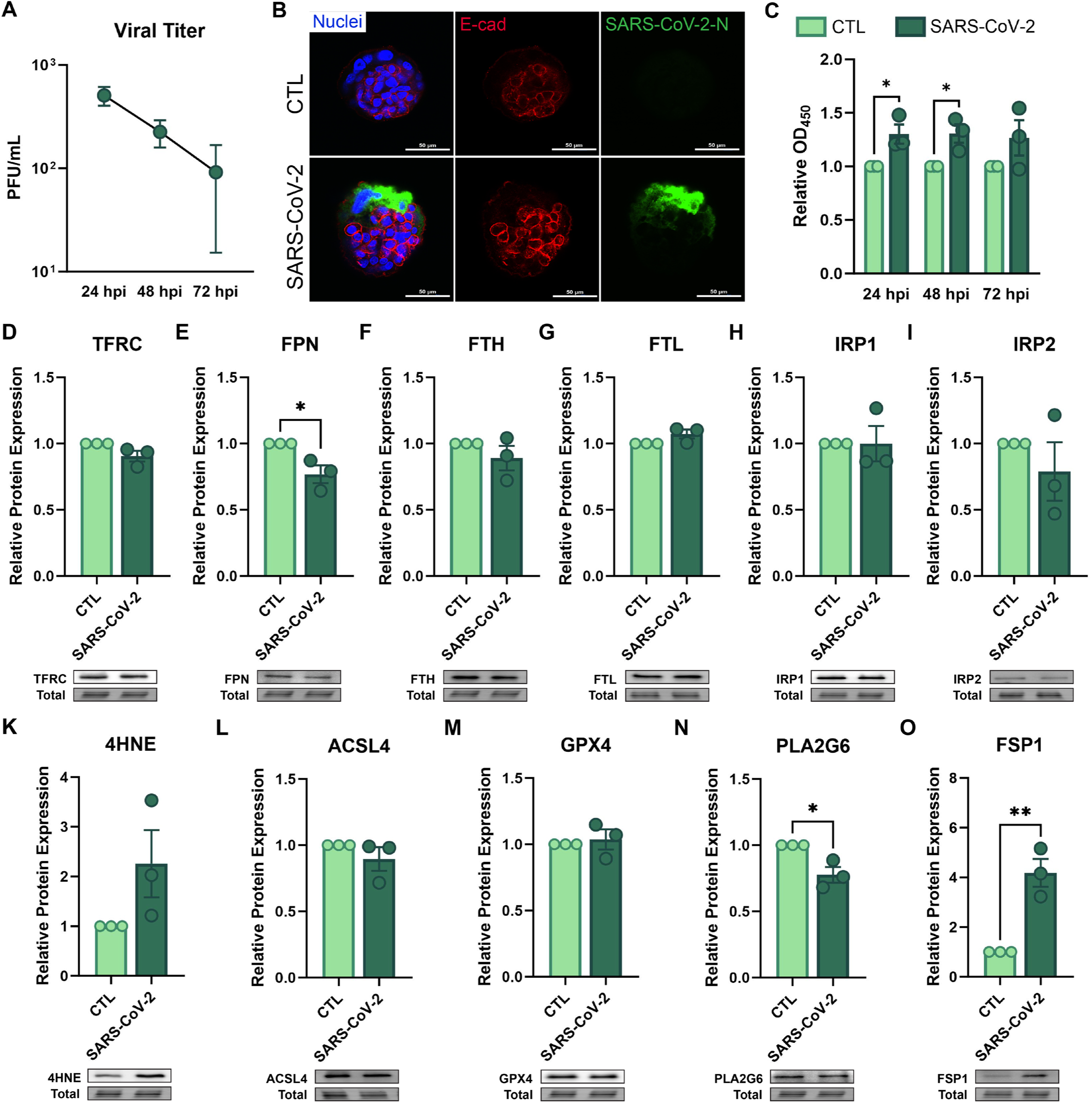
STB^OUT^ SC-TOs infected with SARS-CoV-2 Delta variant exhibit activation of ferroptosis and increased cell death. **(A)** Viral titers of SARS-CoV-2 Delta variant in supernatants from STB^OUT^ SC-TOs infected over 72 hours with an MOI of 10, as determined via plaque assay. **(B)** Representative image of STB^OUT^ SC-TO infected with SARS-CoV-2, as stained with SARS-CoV-2 Nucleocapsid (N, green). Inner CTBs are stained with E-cad (red). **(C)** Cytotoxicity of infected SC-TOs over 72 hpi, as measured by LDH release into supernatant. **(D-O)** Protein expression of iron transport and storage proteins and ferroptosis markers in SARS-CoV-2-infected STB^OUT^ SC-TOs at 72 hpi, as determined by western blotting. Data represents average of n=3 independent replicates per group ± SEM. Significance determined by Student’s unpaired t-test with *p<0.05, **p<0.01.

Similarly to infected JEG-3 cells, SARS-CoV-2 Delta variant increased 4HNE levels, albeit not significantly due to high variability (**Figure 6K**). ACSL4 expression was not significantly affected (**Figure 6L**). In contrast to JEG-3 cells, infected STB^OUT^ organoids exhibited no change in GPX4 expression, but significantly decreased PLA2G6 expression at 72 hpi (**Figure 6M,N**). While this could reflect potential differences between regulation of ferroptosis between JEG-3 and SC-TOs, the discrepancy may also be explained by different time-dependent effects of infection between the two models. Either way, reduced PLA2G6, which protects trophoblasts from ferroptosis,^64^ combined with reduced FPN expression, increased 4HNE levels, and increased cell death in infected organoids, further supports our conclusion that SARS-CoV-2 Delta variant activates trophoblast ferroptosis. However, similar to JEG-3 cells, levels of the ferroptosis inhibitor FSP1 were significantly increased by SARS-CoV-2 (**Figure 6O**), suggesting a potential defense mechanism.

## Discussion

Disrupted cellular iron distribution and activation of ferroptosis have been widely implicated in various non-placental tissues following SARS-CoV-2 infection. In the context of pregnancy, two reports have described increased expression of long-chain-fatty-acid-CoA ligase 4 (ACSL4) in placentas from SARS-CoV-2-infected pregnancies using immunohistochemistry^52,53,65^, suggesting activation of lipid peroxidation pathways upstream of ferroptosis. However, ferroptosis is a multifactorial form of regulated cell death defined by coordinated alterations in iron handling, lipid metabolism, and antioxidant defense, rather than a single molecular executor.^59^ As such, no study has comprehensively examined the impact of SARS-CoV-2 infection on iron dysregulation and ferroptosis in the placenta. Moreover, ferroptosis during viral infection can exert context-dependent effects, functioning either as a host defense mechanism to restrict viral replication, or conversely as a pathway to facilitate infection.^66^ This raises the possibility that ferroptosis may exert distinct, tissue-specific effects at barrier sites such as the placenta. Here, we demonstrate that SARS-CoV-2 infection during pregnancy is associated with impaired placental iron efflux, trophoblast iron accumulation, and activation of ferroptosis signaling. While these changes may contribute to placental dysfunction and adverse pregnancy outcomes associated with COVID-19 during pregnancy, our data also indicate that ferroptosis activation limits viral spread in trophoblasts, consistent with an antiviral role in the placenta. These findings reveal a previously unappreciated trade-off between iron-dependent antiviral defense at the maternal-fetal interface and placental homeostasis.

We observed increased linear deposits of Prussian Blue-stained iron in placentas from SARS-CoV-2-exposed pregnancies, corroborating prior reports that iron levels are elevated in the placenta during COVID-19.^27^ Iron sequestration in exposed placentas was accompanied by reduced ferritin levels in umbilical cord blood, suggesting impaired iron transport to the fetus. Indeed, neonates exposed to SARS-CoV-2 in utero experience increased incidences of anemia.^50,51^ Since iron plays important roles in fetal growth and development, infection-mediated disruption of placental iron handling may help explain these neonatal outcomes, although direct causality remains to be established. The localization of linear iron deposits along the basolateral trophoblast membrane of SARS-CoV-2-exposed placentas, a pattern typically observed in early gestation placentas where iron is stored to meet later fetal demands,^67^ is of particular note. Persistence of these deposits into later pregnancy is associated with fetal pathology, particularly intrauterine growth restriction^67^ and early-onset preeclampsia,^68^ supporting a link between sustained placental iron sequestration and adverse pregnancy outcomes.

Mechanistically, we found that SARS-CoV-2 infection alters expression and localization of ferroportin (FPN), the sole mammalian iron efflux transporter and the primary route for iron transfer from trophoblasts to the fetus. Both JEG-3 cells and STB^OUT^ SC-TOs exhibited reduced FPN expression during active infection with the SARS-CoV-2 Delta variant. Because FPN is the only route through which iron can exit trophoblasts, its downregulation during active infection likely promotes intracellular iron retention. In contrast, FPN protein levels were not significantly altered in SARS-CoV-2-exposed human placentas at term, supporting the idea that this response is acute and restricted to active infection. However, SARS-CoV-2-exposed human placentas displayed a striking alteration in FPN localization at term, with FPN detected on both basolateral and microvillous membranes, an arrangement not previously reported in placental tissue. Apical localization of FPN has been described in airway epithelia, where it is thought to facilitate iron detoxification,^69^ suggesting that altered FPN localization in SARS-CoV-2-exposed placentas may represent a compensatory response to excess intracellular iron. Whether this response is adaptive or ultimately maladaptive warrants further investigation. To assess whether altered iron uptake contributes to placental iron accumulation, we further examined the impact of infection on expression and localization of TFRC, which brings iron into the placenta from maternal circulation. Although SARS-CoV-2 has been reported to use TFRC as an alternative entry receptor^70,71^ and TFRC expression is upregulated in lung tissue of infected monkeys and mice,^70^ we did not observe any change in expression of TFRC in either human placentas or SARS-CoV-2-infected trophoblast models. This suggests that increased placental iron uptake is unlikely to drive iron accumulation during infection.

As for functional consequences of infection-induced iron dysregulation, our results in both trophoblast cells and STB^OUT^ SC-TOs indicate that infection-mediated accumulation of iron, together with decreased expression of the ferroptosis inhibitors GPX4 and/or PLA2G6, promotes activation of ferroptosis signaling in trophoblasts. In contrast, despite evidence of iron accumulation and reduced GPX4 expression, we did not detect robust lipid peroxidation in SARS-CoV-2-exposed placentas. However, multiple studies with larger cohorts who were infected later in gestation have reported increased placental lipid peroxidation,^27,72^ supporting the idea that ferroptosis is most pronounced during active infection and may partially resolve by term if infection occurs in early pregnancy. Ferroptosis has been implicated in various pregnancy complications including preeclampsia, suggesting that transient activation of this pathway during SARS-CoV-2 infection could contribute to placental injury and adverse pregnancy outcomes. Supporting this link, elevated lipid peroxide levels in maternal and neonatal sera following exposure to SARS-CoV-2 during pregnancy^73^ have been correlated with placental vascular pathologies,^74^ and vaccination has been associated with reduced placental lipid peroxidation following infection, suggesting mitigation of ferroptotic injury.^72^ Moreover, our results in JEG-3 cells and SC-TOs suggest that relatively low viral titers are sufficient to cause widespread changes in iron handling and ferroptosis markers, indicative of a large bystander effect. Similar observations have been made during Zika virus infection, where lipid metabolism is altered even in nearby, uninfected trophoblasts,^75^ and could be explained by the fact that ferroptosis can transmit through neighboring cells via plasma membrane contacts.^76^ Moreover, virus-infected cells can exchange cytoplasmic material through additional channels including tunneling nanotubes^77^ and extracellular vesicles,^26,78^ suggesting additional routes through which ferroptotic signals and products could be transmitted between nearby cells. Indeed, iron and lipid peroxides have been identified in placenta-derived small extracellular vesicles during preeclampsia, another condition associated with ferroptosis activation.^79^ While these routes of intercellular communication warrant further investigation in the context of SARS-CoV-2 infection, they provide possible explanations for widespread changes in placental pathology and function, even during or after low levels of placental infection.

Despite its association with placental dysfunction, our results also suggest that ferroptosis activation could be beneficial for limiting viral dissemination in the placenta. Indeed, blocking activity of FSP1, a ferroptosis inhibitor, or activating sub-lethal ferroptosis signaling with RSL3, both reduced viral titers of SARS-CoV-2 Delta variant in infected trophoblasts. This provides a mechanistic explanation for the low levels of SARS-CoV-2 replication typically observed in the placenta and the rarity of vertical transmission.^17–20^ This protective role contrasts with findings in non-placental tissues, where ferroptosis has been shown to promote SARS-CoV-2 replication,^80,81^ prompting investigators to propose ferroptosis inhibition as a therapeutic strategy.^82,83^ However, our results suggest that regulation of SARS-CoV-2 replication by ferroptosis is tissue-specific, raising the possibility that indiscriminate systemic modulation of ferroptosis during pregnancy could have unintended consequences for placental viral control.

A significant strength of our study is the use of STB^OUT^ SC-TOs, which more closely recapitulate placental architecture at the maternal-fetal interface, to model active infection with the SARS-CoV-2 Delta variant. To our knowledge, only two other studies have used 3D trophoblast organoid models to study live infection with SARS-CoV-2. These studies, which include one of our own, used SC-TOs with the STB^IN^ configuration and reported extremely limited infection of only a handful of cells.^44,45^ In contrast, STBs differentiated from hTSCs in 2-dimension are highly susceptible to infection with SARS-CoV-2,^45,63^ likely owing to their higher expression of ACE2.^15,45,63^ We hypothesized that exposing the ACE2-expressing STB layer to culture media by generating 3D STB^OUT^ organoids would increase their susceptibility to SARS-CoV-2 infection. Indeed, we observed robust infection of the outer ACE2-expressing STB layer with the SARS-CoV-2 Delta variant, thereby providing a physiologically relevant 3D platform to study placental SARS-CoV-2 infection.

Our study is not without limitations. The number of human placentas examined was limited and may have hindered our ability to detect statistically significant differences. While our mechanistic studies in JEG-3 cells and STB^OUT^ SC-TOs further support observations derived from human samples, we only examined the impact of the SARS-CoV-2 Delta variant. Because ferroptosis represents a host-encoded response to iron dysregulation and oxidative stress, it is likely to reflect a conserved placental response rather than a variant-specific phenomenon, although this warrants direct testing. Lastly, our studies were conducted in the absence of vaccination, and therefore we cannot draw any conclusions as to whether vaccination may prevent or alleviate the observed changes. Nevertheless, our results provide new mechanistic insight into how SARS-CoV-2 disrupts placental iron homeostasis and activates ferroptosis while simultaneously limiting viral dissemination.

In summary, we identify ferroptosis as a dual-function placental response to SARS-CoV-2 infection that restricts viral replication at the cost of altered iron handling and placental homeostasis. Persistence of infection-induced iron dysregulation beyond viral clearance may contribute to adverse pregnancy outcomes, highlighting the placenta as a site where antiviral defense and developmental demands intersect. Together, these findings underscore the importance of studying host-pathogen interactions within the unique immunometabolic context of pregnancy.

## Methods

### Human Samples

Human placental villous tissue and umbilical cord blood samples were derived from a pre-existing cohort of patient samples collected for the Safety, Testing/Transmission, and Outcomes in pregnancy with COVID-19 (STOP-COVID-19) study at Washington University in St. Louis (IRB approval 202012075).^85^ Samples were collected from consented patients who delivered at Barnes-Jewish Hospital in St. Louis, Missouri between December 2021 and July 2022. Patients were monitored for SARS-CoV-2 infection at enrollment and throughout pregnancy via polymerase chain reaction and/or antigen testing. Patients with any lab-confirmed positive SARS-CoV-2 test during pregnancy were categorized as SARS-CoV-2-exposed (SARS-CoV-2(+), n=10), whereas those negative for SARS-CoV-2 testing during pregnancy were labelled unexposed (SARS-CoV-2(-), n=10).^85^ All individuals included in the present study were unvaccinated against SARS-CoV-2. Exclusion criteria for this patient subset included gestational hypertension, preeclampsia, gestational diabetes, and preterm birth. Placental villous tissue and umbilical cord blood samples were collected at delivery, which did not necessarily coincide with time of active infection.

### Prussian Blue Staining in Human Placentas

Iron deposits were detected in paraffin-embedded placental tissue sections (5 µm) by Prussian Blue Staining with an Iron Stain Kit (Abcam, ab150674) as per manufacturer’s instructions. Briefly, sections were deparaffinized and rehydrated before incubation in a working iron stain solution (potassium ferrocyanide and hydrochloric acid) for 3 minutes. Sections were rinsed thoroughly in distilled water and counterstained in Nuclear Fast Red Solution for 5 minutes. After 4 changes in distilled water, sections were dehydrated and coverslipped with Permount Mounting Medium (Electron Microscopy Solutions, 17986-01). Slides were imaged with a Panoramic 1000 Digital Slide Scanner (3DHistech).

### Quantification of Ferritin in Umbilical Cord Blood

Levels of ferritin were quantified in umbilical cord blood via Luminex. Luminex analysis was performed by Baylor College of Medicine’s Proteomics Core.

### Quantification of Glutathione in Human Placental Tissue

Levels of total (GSSG+GSH) and reduced (GSH) glutathione in frozen placental villous tissue were quantified using a colorimetric GSH+GSSG/GSH Assay Kit (Abcam, ab239709). Approximately 100 mg of tissue was homogenized in Glutathione Reaction Buffer followed by addition of 5-sulfosalicylic acid to remove proteins and peptides. Homogenates were centrifuged at 8000 *g* for 10 mins and the supernatant was used for analysis as per manufacturer’s instructions.

### Quantification of Malondialdehyde in Human Placental Tissue

Malondialdehyde (MDA) levels in frozen placental villous tissue were measured using a colorimetric Lipid Peroxidation (MDA) Assay Kit (Abcam, ab118970). Approximately 25 mg of tissue was homogenized in MDA Lysis Buffer containing butylated hydroxytoluene prior to centrifugation at 13,000 *g* for 10 minutes. The resulting supernatant was used for analysis according to manufacturer’s instructions.

### Cell Culture

The human trophoblast cell line JEG-3 (ATCC HTB-36) was cultured in DMEM/F-12 (Gibco, 11330032) supplemented with 10% Fetal Bovine Serum (FBS) (Gibco, 16140071). The CT30 hTSC line^86^ was a gift from Dr. Thorold W. Theunissen from Washington University in St. Louis. hTSCs were cultured in plates coated with Biolaminin 521 LN (BioLamina, LN521). Media used for culturing hTSCs as cytotrophoblasts (iCTB media) was adapted from Bai *et al.*^87^ with modifications: Advanced DMEM/F12 supplemented with 1X N2, 2X B27, GlutaMax, 0.1 mM 2-mercaptoethanol, 0.05% bovine serum albumin (BSA), 1% knockout serum replacement (KOSR), 0.5% penicillin-streptomycin, 2 uM CHIR99021, 1 uM SB431542, 0.5 uM A83-01, 5 uM Y-27632, 780 uM valproic acid, 50 ng/mL FGF2, and 50 ng/mL EGF. Both cell lines were cultured at 37°C and 5% CO_2_.

### Generation of 3D STB^OUT^ SC-TOs

hTSCs at 80% confluency were used to seed SC-TOs in Matrigel droplets as previously described.^44^ SC-TOs were propagated in Matrigel in 24-well plates with trophoblast organoid medium (TOM)^44^ for three days at 37°C and 5% CO_2_. To generate STB^OUT^ SC-TOs, organoids were removed from Matrigel on Day 3 by trituration with a pipette tip and three cycles of brief centrifugation and washing in Advanced DMEM/F-12. Wells of an AggreWell 400 24-well plate (STEMCELL Technologies, 34415) were treated with Accutase (STEMCELL Technologies, 07920) for 20 minutes at 37°C and washed once with PBS and once with Advanced DMEM prior to organoid seeding. Organoids released from Matrigel were resuspended in TOM complete media and added to the treated AggreWell plate, with organoids derived from one Matrigel dome split between two wells of the AggreWell plate. STB^OUT^ organoids were then maintained statically at 37°C and 5% CO_2_ until infection with SARS-CoV-2 on Day 6.

To determine efficiency of STB^OUT^ production, organoids were collected for immunofluorescence on Days 4, 6, and 8 and stained with markers for STB (SDC-1) and CTB (E-cad) layers. Approximate outer coverage of the STB layer was determined qualitatively by observing cross sections of organoids taken from each timepoint and assigning them to categories of 0-25%, 25-50%, 50-75%, and >75% outer STB coverage.

### Infection of Cells and Organoids with SARS-CoV-2

Stocks of the SARS-CoV-2 Delta strain (B.1.617.2) were prepared in Vero cells (ATCC CCL-81) under Biosafety Level 3 conditions at Baylor College of Medicine. The virus was a donation from Dr. Pedro A Piedra. Viral titers were determined using a standard plaque assay in Vero cells.

Infection of JEG-3 cells and STB^OUT^ organoids were originally performed under BSL-3 conditions, and then under enhanced BSL-2 conditions following the reclassification of SARS-CoV-2’s containment level. JEG-3 cells were seeded in 6-well plates for 24 hours prior to infection at a multiplicity of infection (MOI) of 1. Inoculums were prepared in DMEM/F12 (Gibco) supplemented with 2% FBS (Gibco). After two hours of incubation with the inoculum, cells were returned to standard media with 10% FBS and incubated for 6-72 hours. For experiments involving small molecule inhibitors/activators or iron supplementation, cells were pre-treated with 10 µM iFSP1, 1 µM RSL3, or 400 µM FAC for one hour prior to infection with SARS-CoV-2. After two hours of inoculation, media containing small molecules was re-added to cultures. Conditioned media was collected after 48 hours for cytotoxicity measurements.

Similar procedures were conducted for infection of STB^OUT^ organoids. Organoids were removed from AggreWell plates, briefly centrifuged to remove culture medium, and resuspended in inoculum containing SARS-CoV-2 at an MOI of 10 prepared in Advanced DMEM/F-12 with no supplements. Organoids in inoculum were returned to the AggreWell plates and incubated for two hours, after which organoids were once again removed from the plate, briefly centrifuged, and resuspended in TOM media for continued culture. Viral titers in cell and organoid cultures were determined by subjecting conditioned media to standard plaque assays in Vero cells. In accordance with BSL-3 and enhanced BSL-2 protocols, virus was inactivated in all samples prior to removal from biosafety cabinets.

### Measurement of Cell Death (LDH Assay)

To measure cell death, LDH levels were quantified in cell and organoid conditioned media using a colorimetric CytoTox 96 Non-Radioactive Cytotoxicity Assay (Promega, G1781). Virus in conditioned media was inactivated through incubation with 1% Triton X-100 for 30 minutes prior to conducting the LDH assay.

### Protein Extraction and Western Blotting

Protein was extracted from frozen placental villous tissue (60 mg) using T-PER Tissue Protein Extraction Reagent (Thermo Scientific, 78510). Tissues were homogenized in T-PER containing phosphatase and protease inhibitors and centrifuged at 10,000 *g* for 5 minutes at 4°C. To extract protein from JEG-3 cells infected with SARS-CoV-2, cells were washed with PBS before lysing with ice-cold Pierce RIPA Lysis Buffer (Thermo Scientific, 89901) containing protease and phosphatase inhibitors on ice for 30 minutes. Organoids were prepared for protein extraction by pelleting, washing three times in PBS containing 0.1% BSA, and snap-freezing in liquid nitrogen. After a freeze-thaw cycle, pellets were resuspended in RIPA Lysis Buffer and passed through a 27-gauge needle three times before incubation on ice for 30 minutes for virus inactivation. Cell and organoid lysates were subsequently sonicated at 20% amplitude for five rounds of 2 seconds on with 8 seconds off. Lysates were centrifuged at 14,000 *g* at 4°C for 15 minutes. Protein was quantified in supernatants from tissue, cells, and organoids using a Pierce BCA Protein Assay Kit (Thermo Scientific, 23225) as per manufacturer’s instructions.

Protein (25 µg) was subjected to SDS-PAGE on 4-20% gradient Mini-PROTEAN TGX stain-free precast gels (Bio-Rad Laboratories, 4561093) and transferred to PVDF membranes at 80 V for two hours. Total protein signals were acquired using stain-free technology with a Bio-Rad Chemidoc system. Membranes were blocked with TBS Intercept Blocking Buffer (LICORbio, 927-60001) for one hour at room temperature before incubation with primary antibodies (listed in Supplementary Table S1) overnight at 4°C. Membranes were subsequently washed five times with Tris-buffered saline containing 0.1% Tween-20 (TBST) before incubation with infrared dye-linked fluorescent secondary antibodies (LICORbio, **Table S1**) for one hour at room temperature. After another series of five washes in TBST, proteins were imaged on a ChemiDoc using infrared detection. Band intensities were analyzed in Image Lab (Bio-Rad) and normalized to total protein.

### Immunofluorescence

Paraffin-embedded placental villous tissue samples were sectioned into 5 µm slices and subjected to deparaffinization and rehydration. Heat-mediated antigen retrieval was performed in 10 mM sodium citrate with 0.05% Tween (pH 6.0) using a pressure cooker for all antigens except Spike protein. Tissue sections were washed in PBS prior to blocking in 5% normal horse serum for one hour at room temperature. Primary antibodies prepared in 5% horse serum were applied to slides overnight at 4°C. Stained slides were washed three times with PBS prior to staining with corresponding Alexa Fluor-conjugated secondary antibodies prepared in 1% horse serum for 1 hour at room temperature. After three washes in PBS, autofluorescence was quenched using the Vector TrueVIEW Autofluorescence Quenching Kit (Vector Laboratories, SP-8400) as per manufacturer’s instructions. Sections were counterstained with Hoechst 33342 (1:10,000; Invitrogen, H3570) for five minutes and coverslipped with VECTASHIELD Vibrance Antifade Mounting Medium (Vector Laboratories, H-1700). Slides were imaged with an ECLIPSE Ti2 confocal microscope (Nikon) within 24 hours of staining.

For immunofluorescence staining of SC-TOs, organoids were fixed with 4% paraformaldehyde for 30 minutes at room temperature and washed three times with 0.1% BSA in PBS. Organoids were blocked, permeabilized, and stained with antibodies (**Table S1**) and Hoechst 33342 during overnight incubations and as described previously.^44^ Organoids were mounted in a fructose glycerol clearing solution suspended between two coverslips separated by CoverWell Incubation Chambers (Grace Bio-Labs, 645401) and imaged with an ECLIPSE Ti2 confocal microscope (Nikon).

### Statistical Analysis

Statistical analysis was conducted in GraphPad Prism 9. For patient demographic information, continuous variables were compared using Mann-Whitney U tests and categorical variables were assessed via Fisher’s exact test. For other analyses, Student’s t-test or Mann-Whitney U tests were used to compare between two groups, depending on normality as determined by Shapiro-Wilk test. For comparisons between more than two groups, one-way ANOVA with Tukey’s multiple comparison test was used. To determine significance between control and infected samples treated with small molecules, two-way ANOVAs with Tukey’s multiple comparison test were used.

## Acknowledgements

This work was supported in part by an NIH/NICHD R01HD091218-04S1 to IUM and JCK and NIH/NIAID R01AI176505 to IUM. ERM was supported by a Postdoctoral Fellowship in Infection and Immunity from Baylor College of Medicine. DK was supported by the Early Career Award Program from Thrasher Research Fund. BRJ was a recipient of an NIH/NIGMS training grant 5T32GM136554-03 and a grant to Baylor College of Medicine from the Howard Hughes Medical Institute through the Gilliam Fellows Program. The work in the Theunissen lab was supported by an NIH Director’s New Innovator Award (DP2GM137418), NIH/NIGMS R35GM153439-01, NIH/NICHD R01HD119277-01, and grants from the Shipley Foundation Program for Innovation in Stem Cell Science and the Edward Mallinekrodt, Jr. Foundation. JCK and ED were also supported by NIH/NIDA R21DA057493-02, NIH/NICHD 1R01HD113199-01, and NIH/NIDA R61DA062321-01. Several figures in this manuscript were generated using BioRender.com.

## Materials Availability

Further information and requests for resources and reagents should be directed to and will be fulfilled by the lead contact, Indira Mysorekar.

---

**Supplementary Table S1:**
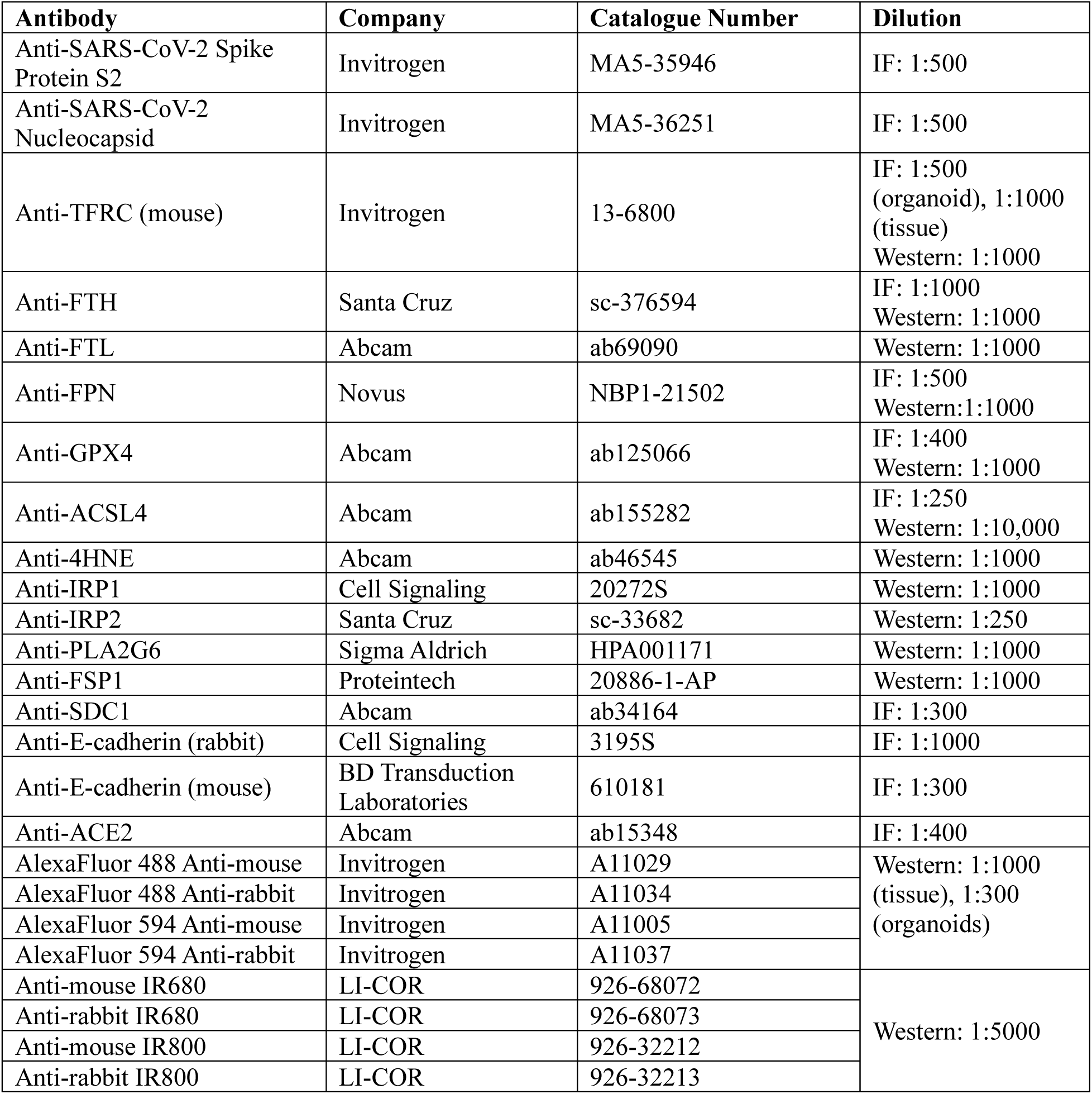
Antibodies used for western blotting and immunofluorescence.

