## Supplementary material for "Trophoblast ferroptosis restricts SARS-CoV-2 spread in the placenta"

**Supplementary Table S1:** Antibodies used for western blotting and immunofluorescence.

| Antibody | Company | Catalogue Number | Dilution |
|---|---|---|---|
| Anti-SARS-CoV-2 Spike Protein S2 | Invitrogen | MA5-35946 | IF: 1:500 |
| Anti-SARS-CoV-2 Nucleocapsid | Invitrogen | MA5-36251 | IF: 1:500 |
| Anti-TFRC (mouse) | Invitrogen | 13-6800 | IF: 1:500 (organoid), 1:1000 (tissue)<br>Western: 1:1000 |
| Anti-FTH | Santa Cruz | sc-376594 | IF: 1:1000<br>Western: 1:1000 |
| Anti-FTL | Abcam | ab69090 | Western: 1:1000 |
| Anti-FPN | Novus | NBP1-21502 | IF: 1:500<br>Western:1:1000 |
| Anti-GPX4 | Abcam | ab125066 | IF: 1:400<br>Western: 1:1000 |
| Anti-ACSL4 | Abcam | ab155282 | IF: 1:250<br>Western: 1:10,000 |
| Anti-4HNE | Abcam | ab46545 | Western: 1:1000 |
| Anti-IRP1 | Cell Signaling | 20272S | Western: 1:1000 |
| Anti-IRP2 | Santa Cruz | sc-33682 | Western: 1:250 |
| Anti-PLA2G6 | Sigma Aldrich | HPA001171 | Western: 1:1000 |
| Anti-FSP1 | Proteintech | 20886-1-AP | Western: 1:1000 |
| Anti-SDC1 | Abcam | ab34164 | IF: 1:300 |
| Anti-E-cadherin (rabbit) | Cell Signaling | 3195S | IF: 1:1000 |
| Anti-E-cadherin (mouse) | BD Transduction Laboratories | 610181 | IF: 1:300 |
| Anti-ACE2 | Abcam | ab15348 | IF: 1:400 |
| AlexaFluor 488 Anti-mouse | Invitrogen | A11029 | Western: 1:1000 (tissue), 1:300 (organoids) |
| AlexaFluor 488 Anti-rabbit | Invitrogen | A11034 | |
| AlexaFluor 594 Anti-mouse | Invitrogen | A11005 | |
| AlexaFluor 594 Anti-rabbit | Invitrogen | A11037 | |
| Anti-mouse IR680 | LI-COR | 926-68072 | Western: 1:5000 |
| Anti-rabbit IR680 | LI-COR | 926-68073 | |
| Anti-mouse IR800 | LI-COR | 926-32212 | |
| Anti-rabbit IR800 | LI-COR | 926-32213 | |
